# Codon usage influences fitness through RNA toxicity

**DOI:** 10.1101/344002

**Authors:** Pragya Mittal, James Brindle, Julie Stephen, Joshua B. Plotkin, Grzegorz Kudla

**Affiliations:** MRC Human Genetics Unit, IGMM, University of Edinburgh, Edinburgh, Scotland, UK; University of Pennsylvania, Philadelphia, USA

## Abstract

Many organisms are subject to selective pressure that gives rise to unequal usage of synonymous codons, known as codon bias. To experimentally dissect the mechanisms of selection on synonymous sites, we expressed several hundred synonymous variants of the GFP gene in *Escherichia coli*, and used quantitative growth and viability assays to estimate bacterial fitness. Unexpectedly, we found many synonymous variants whose expression was toxic to *E. coli*. Unlike previously studied effects of synonymous mutations, the effect that we discovered is independent of translation, but it depends on the production of toxic mRNA molecules. We identified RNA sequence determinants of toxicity, and evolved suppressor strains that can tolerate the expression of toxic GFP variants. Genome sequencing of these suppressor strains revealed a cluster of promoter mutations that prevented toxicity by reducing mRNA levels. We conclude that translation-independent RNA toxicity is a previously unrecognized obstacle in bacterial gene expression.

**Significance statement:** Synonymous mutations in genes do not change protein sequence, but they may affect gene expression and cellular function. Here we describe an unexpected toxic effect of synonymous mutations in *Escherichia coli*, with potentially large implications for bacterial physiology and evolution. Unlike previously studied effects of synonymous mutations, the effect that we discovered is independent of translation, but it depends on the production of toxic mRNA molecules. We hypothesize that the mechanism we identified influences the evolution of endogenous genes in bacteria, by imposing selective constraints on synonymous mutations that arise in the genome. Of interest for biotechnology and synthetic biology, we identify bacterial strains and growth conditions that alleviate RNA toxicity, thus allowing efficient overexpression of heterologous proteins.

## Main text

Although synonymous mutations do not change the encoded protein sequence, they cause a broad range of molecular phenotypes, including changes of transcription ^1^, translation initiation^2, 3^, translation elongation^4^, translation accuracy^5, 6^, RNA stability^7^, and splicing^8^. As a result, synonymous mutations are under subtle but non-negligible selective pressure, which manifests itself in the unequal usage of synonymous codons across genes and genomes^9^^-^^11^. Several recent experiments directly measured the effects of synonymous mutations on fitness in bacteria^2, 12-17^. It has been commonly assumed that fitness depends primarily on the efficiency, accuracy, and yield of translation. Here we show that in the context of heterologous gene expression in *E. coli*, large effects of synonymous mutations on fitness are translation-independent, and are mediated by RNA toxicity.

To study the effects of synonymous mutations on bacterial fitness, we used an IPTG-inducible, bacteriophage T7 polymerase-driven plasmid to express a collection of synonymous variants of the GFP gene^2^ in *E. coli* BL21-Gold(DE3) (henceforth referred to as BL21) cells (see Methods). Without IPTG induction, there were no discernible differences in growth between strains (Figure 1A). When induced with IPTG, the growth rate of GFP-producing strains was reduced, consistent with the metabolic burden conferred by heterologous gene expression. The growth phenotype varied remarkably between strains expressing different synonymous variants of GFP (Figure 1B, Supp Figure 1). “Slow” variants caused a long lag phase post-induction, indicating that at this stage the cells either stopped growing or died, while ‘fast’ variants showed growth rates closer to non-induced cells. Several hours after induction, the slow variants appeared to resume growth (Figure 1B): we found that this was related to the emergence of suppressor strains that could tolerate the expression of these variants (Supp Figure 1D, and see below).

**Figure 1.**
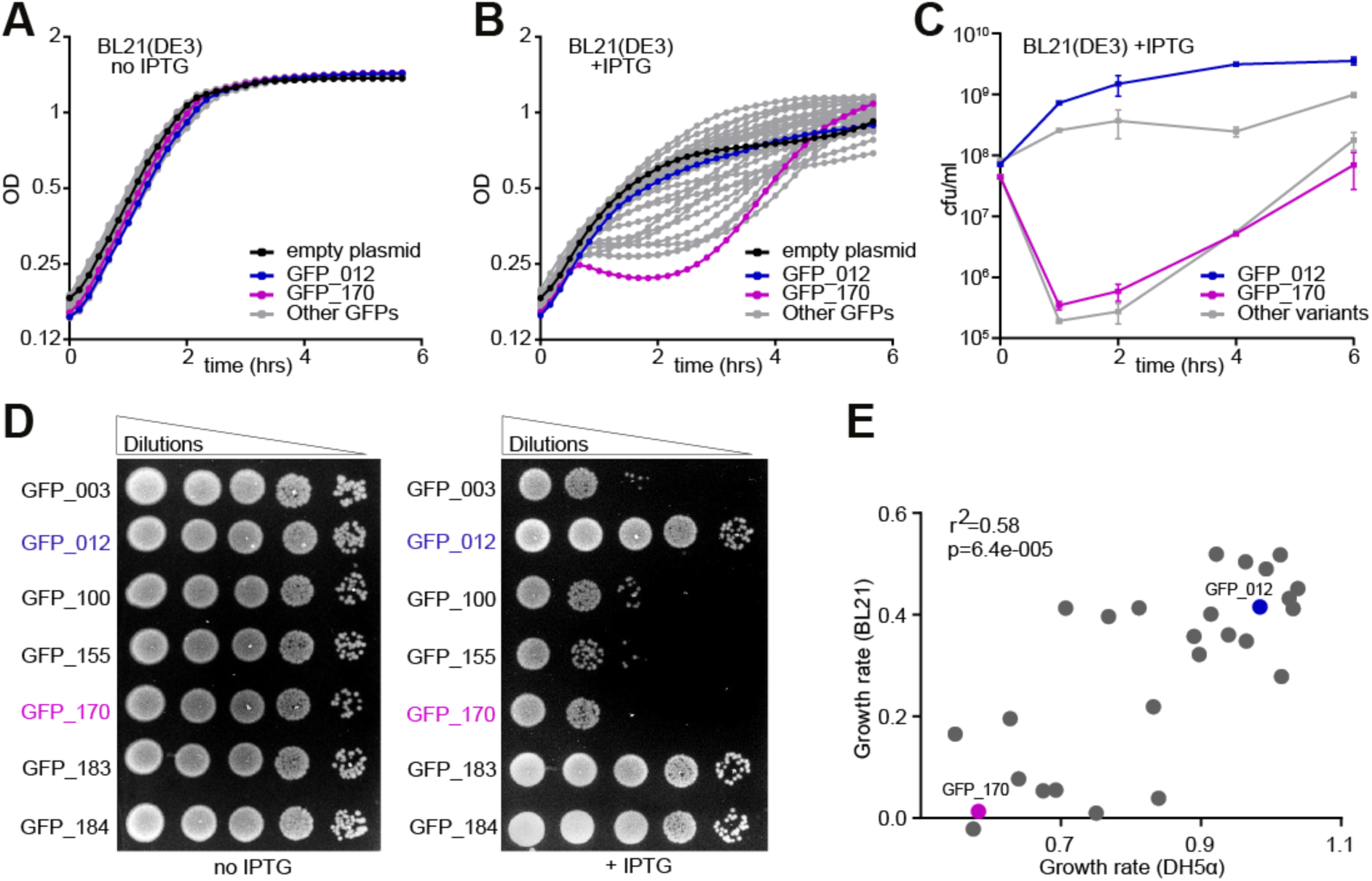
GFP variants are toxic in *E. coli*. **(A-B)** Growth curves of BL21 *E. coli* cells, non-induced (A) or induced with 1 mM IPTG at t=0h (B). Cells carrying GFP_012 (non-toxic variant, blue), GFP_170 (toxic variant, magenta), pGK8 (empty vector control, black) and 29 other variants (grey) are shown. Each curve represents an average of 9 replicates (3 biological × 3 technical). OD, optical density. **(C)** Numbers of colony forming units (cfu)/ml at specified time points after induction with 1 mM IPTG. Data points represent averages of 4 replicates, +/- SEM. **(D)** Semi-quantitative estimation of BL21 cell viability by spot assay. **(E)** Estimated growth rates of cells expressing GFP variants in DH5α and BL21 strains (averages of at least 6 replicates).

We quantified cell viability post-induction by assessing the colony-forming ability of cells (Figure 1C). Fast variants showed the expected increase in cell numbers post-induction, but slow variants caused a 1000-fold decrease in viable cell numbers. Similarly, spotting of non-induced cells onto LB plates with IPTG showed that the slow variants formed markedly fewer colonies than fast variants (Figure 1D). Microscopic analysis of slow variants showed decrease in cell number, growth arrest and in some cases massive cell death following IPTG induction. In the case of fast variants we observed normal increase in cell numbers and negligible cell death after induction (Supp Figure 2). These results indicate that certain synonymous variants of GFP cause significant growth defects when overexpressed in *E. coli* cells, and we will henceforth refer to these variants as “toxic”.

To test if toxicity was specific to T7 promoter-driven overexpression, we analysed growth phenotypes following the expression of a subset of GFP variants using a bacterial polymerase (*trp*/*lac*) promoter system (Methods). Although the growth phenotypes measured with bacterial promoter constructs were not as dramatic as with T7-based constructs, presumably because of lower GFP expression levels, growth rates with both types of promoters were correlated with each other (Figure 1E). Interestingly, toxicity increased at high temperature, and decreased at low temperature (Supp Figure 1C). Taken together, these results indicate that the toxic GFP variants cause growth defects in two different *E. coli* strains, with two types of promoters, possibly through a common mechanism.

To understand if toxicity depends on the process of translation, we selected several toxic and nontoxic variants of GFP and mutated their Shine-Dalgarno (SD) sequences from GAAGGA to TTCTCT to prevent ribosome binding and block translation initiation. As expected, mutation of SD sequences completely inhibited the production of functional GFP protein from all tested constructs (Figure 2A). To our surprise, GFP variants without SD sequences remained toxic, and their effects on growth were indistinguishable from variants with a functional SD sequence (Figure 2B). Western blot analysis confirmed that mutation of the SD sequences ablates GFP expression (Supp Figure 3). We considered the possibility that a cryptic SD element within the coding region allowed translation of a truncated fragment of GFP, which would be consistent with loss of GFP fluorescence and translation-dependent toxicity. However, analysis of the coding regions with the RBS Calculator^18^ revealed no strong SD consensus sequences. These results raise the possibility that toxicity might arise at the RNA level, rather than at translation or protein level.

**Figure 2.**
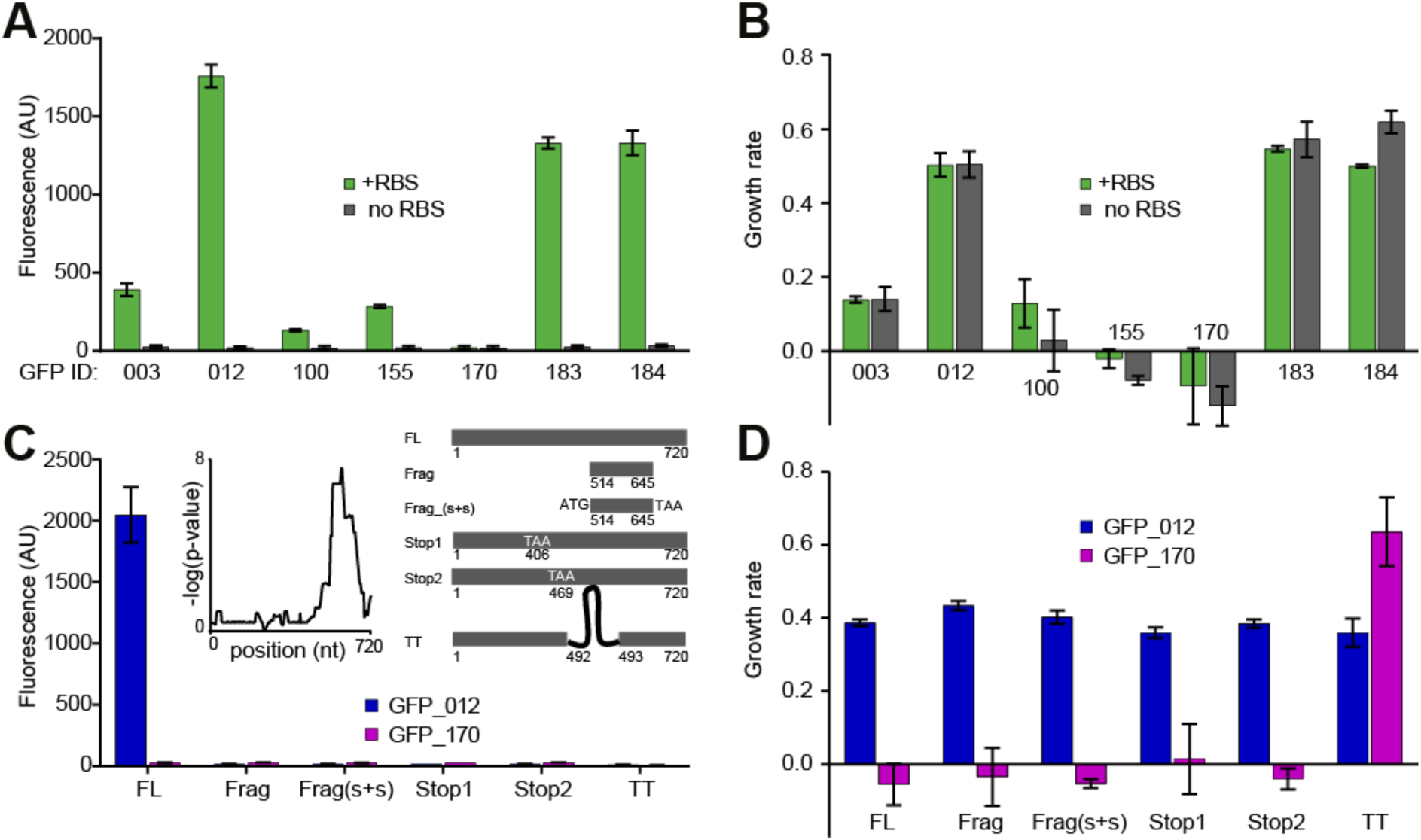
Toxicity of GFP variants is independent of translation. **(A-B)** Fluorescence (A) and growth rate (B) of BL21 cells expressing GFP variants with functional and non-functional ribosome-binding sites (RBS). **(C-D)** Fluorescence (C) and growth rate (D) of cells expressing full-length GFP variants, truncated variants, and variants containing internal stop codons or transcription terminators. Inset in (C) shows location of toxic sequence element in GFP_170 which was calculated based on an analysis of growth rates of 36 shuffled constructs. The Y-axis shows the statistical significance of the association of particular positions with slow growth. Variants derived from non-toxic GFP_012 are shown in blue, and variants derived from toxic GFP_170 are shown in magenta. Full-length constructs, truncated constructs and constructs with internal stop codons have similar growth rates, suggesting that the element of toxicity resides within the truncated fragment and that the mechanism of toxicity is independent of translation. FL, full-length construct; TT, T7 transcription terminator. All data are averages of 9 replicates, +/- SEM.

To identify sequence elements required for toxicity, we selected one of the toxic variants (GFP_170), and a nontoxic variant (GFP_012), and performed DNA shuffling^19^ to generate constructs that consisted of random fragments of GFP_170 and GFP_012. All the shuffled and non-shuffled constructs we generated encoded the same GFP protein sequence. Analysis of growth rate phenotypes of these shuffled constructs revealed a fragment near the 3′ end of the GFP_170 coding sequence (nt 514-645) that was sufficient to elicit the toxic phenotype (Figure 2C, Supp Figure 4A, B). Some mutations outside of the toxic region partially improved fitness, which might be explained by interactions of the RNA secondary structure between the toxic region and the mutated regions. The GFP_170 mRNA is predicted to have a very low translation initiation rate, due to strong RNA secondary structure near the mRNA 5′ end^2^. Nevertheless, replacement of the strongly structured 5′ region with an unstructured fragment did not affect toxicity (Supp Figure 4A, B).

The above results led us to hypothesize that the toxicity associated with GFP expression was independent of translation, but depended on the presence of a specific fragment of RNA. To test this hypothesis, we performed growth rate measurements with a series of constructs. First, we isolated the 132-nt toxic region identified in the DNA shuffling experiment, and expressed it on its own, with or without start and stop codons. The expression of the 132-nt fragment of GFP_170 was sufficient for toxicity, whereas the corresponding fragment of GFP_012 did not cause toxicity. The effect of the 132-nt fragments on growth did not depend on the presence of translation start and stop codons (Figures 2C, D), the fragments contained no cryptic translation initiation signals, and FLAG tag fusions showed no detectable protein expression from the GFP_170 fragment in any of the three reading frames (Supp Figure 3B). Second, we introduced stop codons upstream of the toxic fragment in the GFP_170 coding sequence, and in the corresponding positions of GFP_012. This placement of stop codons ensures that ribosomes terminate translation before reaching the putative toxic region of the RNA, while still allowing a full-length transcript to be produced. As expected, internal stop codons abrogated GFP protein production (Figure 2C), but despite the presence of premature stop codons, GFP_170_Stop still caused toxicity to bacterial cells while GFP_012_Stop remained non-toxic (Figure 2D). To remove possible out-of-frame translation, we inserted stop codons into GFP_170 in all three frames, before and after the toxic region, and toxicity remained the same in all cases (Supp Figure 4C). Third, we introduced an efficient synthetic T7 transcription terminator^20^ upstream of the toxic region in GFP_170 and in the corresponding location in GFP_012. Notably, we found that both variants with internal transcription terminators became nontoxic, and GFP_170_TT grew slightly faster than GFP_012_TT (Figure 2D). The GFP_170 fragment also caused toxicity when fused to FLAG tags (in any of the three reading frames), and when fused to fluorescent protein mKate2, it caused toxicity and reduced expression of mKate2 by 50-fold (Supp Figure 4D, E, F). Overall, these data suggest that toxicity is caused by the RNA itself, rather than the process of translation or by the protein produced.

To investigate the sequence determinants of RNA-mediated toxicity, we measured the growth phenotypes of single synonymous mutations within the 132-nt region of GFP_170. Close to half of these mutations reduced or abolished the toxic phenotype, whereas the remaining mutations had no effect (Figure 3A). There was no clear relationship between the position of mutations within the region and their effect on growth, nor was there any relationship between the type of nucleotide introduced and growth. RNA toxicity associated with triplet repeats has been described in Eukaryotes^21^, but we found no triplet repeats in the toxic GFP mRNAs. Consistent with our observation that the toxic effect does not require translation, codon adaptation index was not associated with toxicity (Figure 3B). RNA folding energy, measured either in the immediate vicinity of each mutation, or for the entire 132-nt mutagenized region, was not correlated with toxicity, and we were unable to identify any RNA structural elements associated with the toxic phenotype (data not shown). We further probed the effects of sets of several mutations within the 132-nt toxic region. 75/98 sets of mutations we introduced within the region reduced or abolished toxicity, whereas 23/98 sets had no effect (Supp Figure 5). In almost all cases, the phenotypes of sets could be deduced from the effects of individual mutations in a simple way: if any mutation in a set abolished toxicity, then the set also did. Four sets did not conform to this rule, indicating potential epistatic interactions between mutations (not shown). Mutations near the 3′ end of the 132-nt fragment had no effect on toxicity, identifying a minimal toxicity-determining region of about a hundred nucleotides that either consists of a single functional element, or it contains multiple elements whose cooperative action causes toxicity.

**Figure 3.**
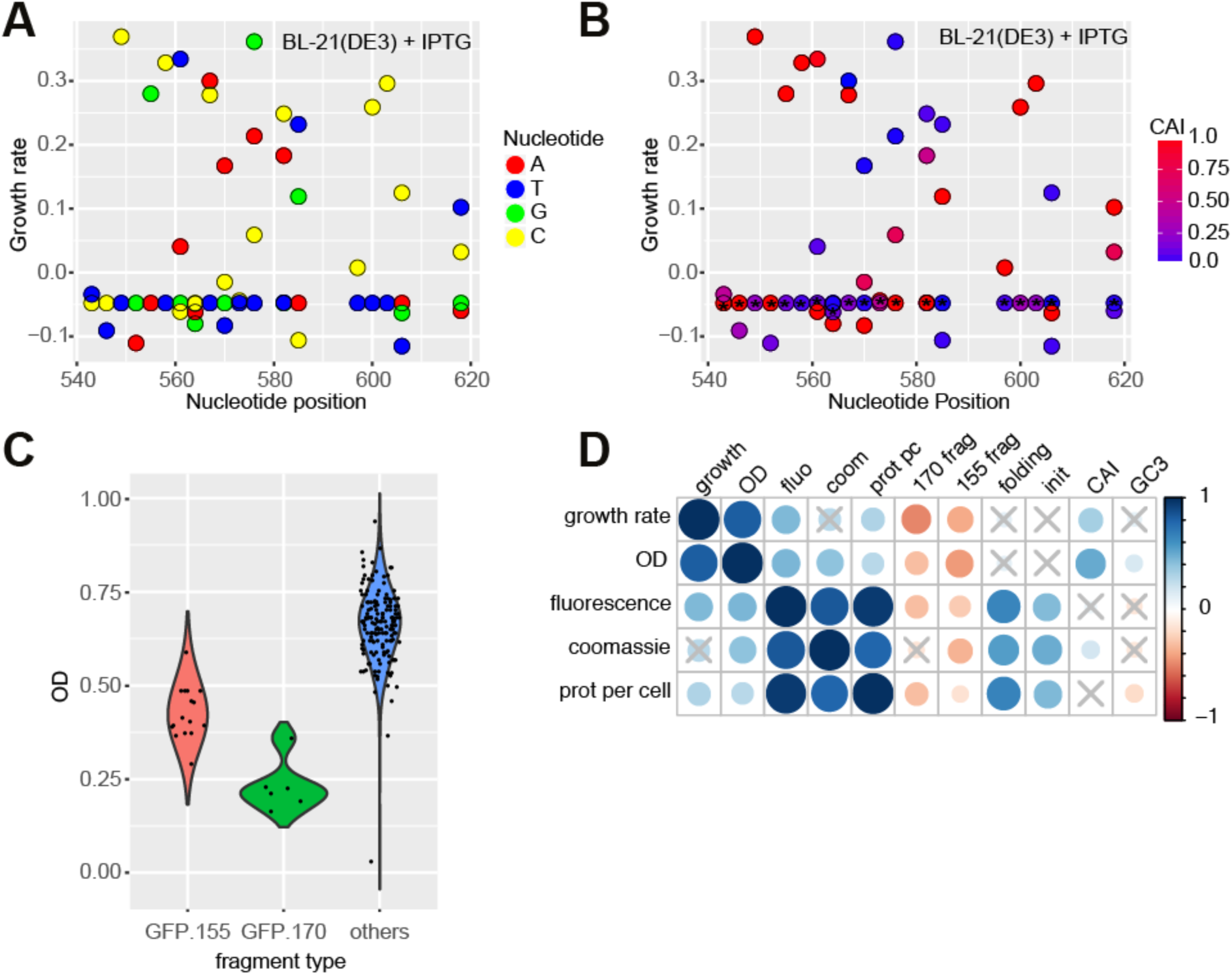
Multiple sequence elements determine RNA-mediated toxicity. **(A)** Growth rates of single synonymous mutants of GFP_170, measured in BL21 strain (averages of 9 replicates). Mutations located throughout the toxic region reduce or abolish toxicity. **(B)** Relationship between Codon Adaptation Index (CAI) and the growth rate of GFP mutants. Asterisk-marked codons represent the original codon in GFP_170. **(C)** Growth estimate (optical density) of BL21 cells expressing GFP variants containing fragments: GFP_155 nt 490-720 (N=16, red), GFP_170 nt 514-645 (N=6, green), and other variants (N= 163, blue). **(D)** Spearman correlation analysis of phenotypes measured in BL21 cells and sequence covariates in a set of 190 GFP variants. The size and colour of circles represents the correlation coefficient; crosses indicate non-significant correlations.

Several recent studies examined the effects of synonymous mutations on fitness in bacteria, either in endogenous genes, or in overexpressed heterologous genes^2, 12-16^. Fitness had been found to correlate with the codon adaptation index (CAI), GC content, RNA folding, protein expression level, a codon ramp near the start codon, and measured or predicted translation initiation rates. We quantified these variables in a set of 190 synonymous variants of GFP, and analysed their impact on fitness. We also considered two candidate toxic RNA fragments (GFP_170, nt 514-645, and GFP_155, nt 490-720), both of which were common to several constructs and appeared to negatively influence fitness (Figures 3C, D). High protein expression was previously shown to correlate with slow growth^14^, whereas we found positive correlations of fitness with total protein yield or protein yield per cell. These correlations presumably reflect reduced protein yields and cell growth after the induction of toxic RNAs. As seen previously, growth rate and optical density were positively correlated with CAI, and GC content was correlated with optical density^2, 16^. However, in a multiple regression analysis aimed to disentangle the effects of these covariates, we found that the presence of candidate toxic RNA fragments predicted slow growth in both BL21 and DH5α cells, whereas CAI and GC3 did not (Methods). This suggests that the apparent correlation of CAI or GC content with fitness, observed in this and previous studies^2, 16^, might result from the confounding effect of toxic RNA fragments (Supp Figure 6A, B). Consistently, an experiment with 22 new, unrelated synonymous GFP constructs spanning a wider range of GC content showed no correlation between GC content and bacterial growth (Supp Figure 6C, D). To further test whether toxicity could be explained by unusually high expression of certain GFP variants, we measured the mRNA abundance of 79 toxic and non-toxic RNAs by Northern blots, and correlated GFP mRNA abundance per cell with OD. Although we observed differences in mRNA abundance, mostly related to mRNA folding^2^, we find no significant correlation between RNA abundance and toxicity (Spearman rho=0.12, p=0.29). Furthermore, we detected no consistent differences in plasmid abundance between toxic and nontoxic variants.

To study the molecular mechanisms of toxicity caused by mRNA overexpression, we aimed to evolve genetic suppressors of this phenotype. We selected several GFP constructs that showed both strong toxicity and moderate or high GFP fluorescence, and plated bacteria containing these constructs on LB agar plates with IPTG and ampicillin. We observed a number of large white colonies that apparently expressed no GFP, and smaller bright green colonies producing high amounts of the GFP protein (Figure 4A). We hypothesized that the green colonies have acquired a genomic mutation that allowed cells to survive while expressing toxic RNAs. To support this, we cured the evolved strains of their respective plasmids and re-transformed the cured strains with the same plasmid. The re-transformed strains readily formed bright green colonies on IPTG+ampicillin plates, and exhibited faster growth rates in IPTG medium compared to the parental strain. This supported our hypothesis that the mutations were located on the chromosome and not the plasmid. We therefore selected 22 evolved strains and the parental strain for genome sequencing, and used the GATK pipeline for calling variants (Methods).

**Figure 4.**
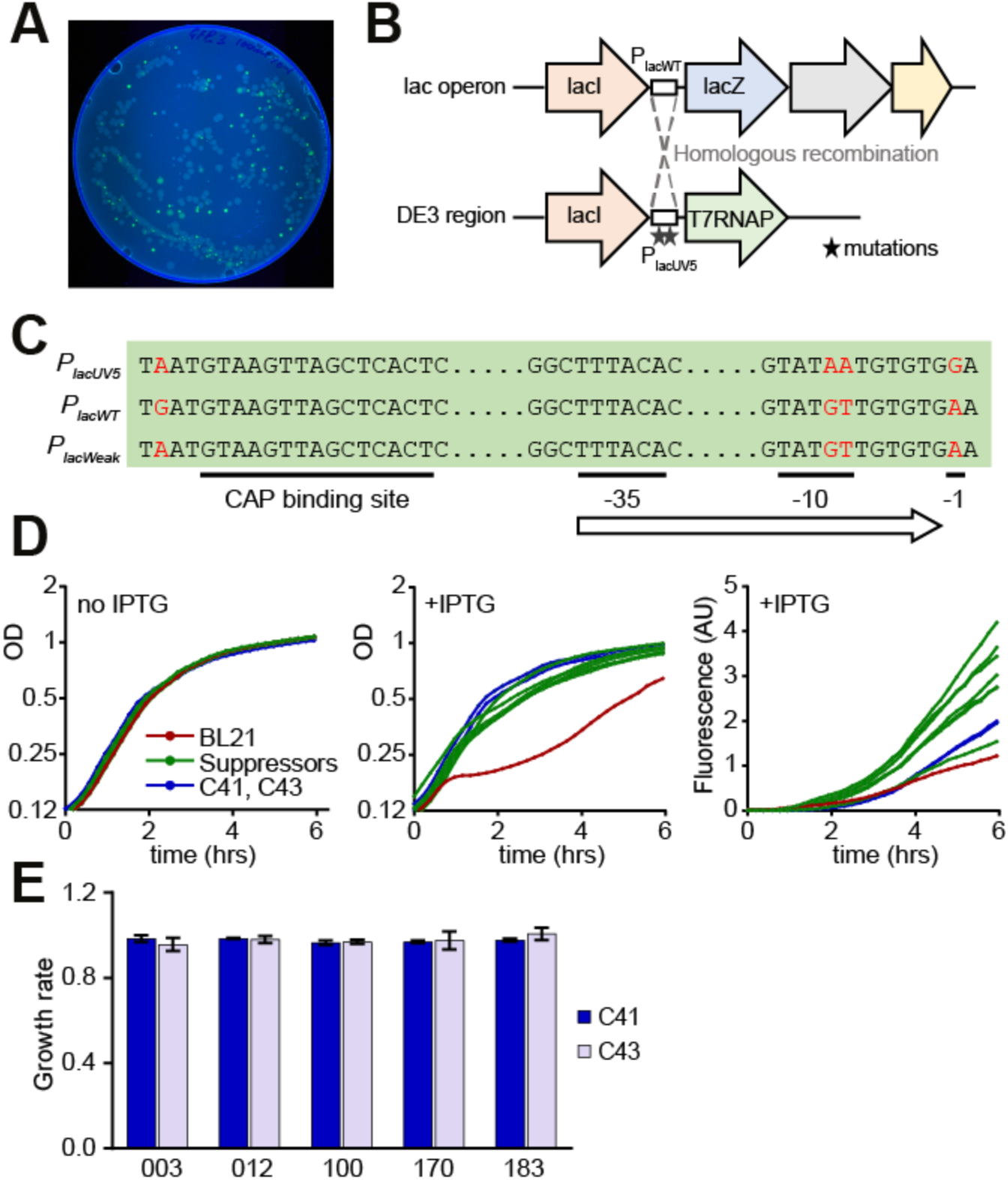
Isolation and characterization of genetic suppressors of toxicity. (**A)** Fluorescence image of LB+Amp+IPTG Petri dish with BL21 cells expressing GFP_003 variant. **(B)** Genetic organization of *lac* and DE3 loci in BL21 cells. Dashed lines indicate homologous recombination between the loci in suppressor strains. **(C)** Sequence variation between the three types of promoters found in the suppressor strains. Substitutions are marked in red. **(D)** Growth curves and fluorescence of strains carrying the GFP_003 variant: parental BL21 strain (red), suppressors strains (N=7, green), C41 and C43 strains (blue). **(E)** Growth rates of C41 and C43 cells expressing several GFP variants. GFP_003, GFP_100 and GFP_170 are toxic in the BL21 strain, GFP_012 and GFP_183 are not. Growth curves are averages of 3 replicates.

In all green suppressor strains, we found a single cluster of mutations in the P*_lac_* promoter of the T7 polymerase gene that explains the suppressor phenotype (Figure 4B, C, Supp Table 1). The parental BL21 strain contains two alleles of the P*_lac_* promoter: the wild-type allele P*_lac_*_WT_ controls the lac operon, and a stronger derivative allele P*_lac_*_UV5_ controls T7 RNA polymerase. In the suppressor strains, recombination between these two loci associates P*_lac_*_WT_ promoter with T7 polymerase, leading to reduced levels of polymerase and presumably to reduced transcription of GFP. The same P*_lac_* promoter mutations were recently observed in the C41(DE3) and C43(DE3) strains of *E. coli* (the ‘Walker strains’), and were responsible for the reduced T7 RNA polymerase expression, high-level recombinant protein production, and improved growth characteristics of those strains^22^^-^^24^. Similar to our suppressor strains, C41(DE3) and C43(DE3) allowed high protein expression of toxic GFP variants, and little toxicity was observed in these strains (Figure 4D). Taken together, these results support our conclusion that high levels of RNA, rather than RNA translation or protein, are responsible for toxicity.

To test whether translation-independent RNA toxicity might affect genes other than GFP, we turned to the *ogcp* gene, which encodes a membrane protein Oxoglutarate-malate transport protein (OGCP) believed to be toxic for *E. coli*. OGCP overexpression was originally used to derive the C41( DE3) strain, now commonly used for recombinant protein expression^22^. As expected, we found that expression of OGCP was toxic to BL21 but not to C41(DE3) cells. In agreement with our observations for GFP, a translation-incompetent variant of OGCP lacking the Shine-Dalgarno sequence was just as toxic to BL21 cells as a translation-competent variant (Supp Figure 7). A translation-competent, codon-optimized variant of OGCP retained toxicity in BL21 cells. These experiments suggest that translation-independent RNA toxicity might be a widespread phenomenon associated with heterologous gene expression in *E. coli*. Heterologous protein expression is known to inhibit growth of *E. coli*. Toxicity is typically attributed to the foreign protein itself, and it is often remedied by lowering expression, reducing growth temperature, or using special strains of *E. coli* such as C41(DE3). Here we demonstrate that the same strategies and strains also prevent toxicity when RNA, rather than protein, is the toxic molecule. We speculate that other cases of toxicity, previously attributed to proteins, may in fact be caused by RNA. Although the molecular mechanisms of RNA toxicity are presently unclear, we identified several GFP and OGCP variants with similar phenotypes, suggesting that the phenomenon may be common. Interestingly, induction of wild-type APE_0230.1 in *E. coli* inhibits growth, but a codon-optimized variant does not inhibit growth despite increased protein yield^25^. In addition, several recent high-throughput studies found unexplained cases of slow growth or toxicity upon the expression of various random sequences in *E. coli*^14, 26, 27^. Our results point to RNA toxicity as a possible cause of these observations.

Our results are relevant to the phenomenon of synonymous site selection in microorganisms. Synonymous mutations can influence fitness directly (in cis), by changing the expression of the gene in which the mutation occurs^12, 13, 15^, or indirectly (in trans), by influencing the global metabolic cost of expression^2, 14, 16, 28^. Experiments with essential bacterial genes predominately uncover cis-effects, most of them mediated by changes of RNA structure or other properties that influence translation yield. For example, mutations in *Salmonella enterica rpsT* downregulated the gene, and could be compensated by additional mutations in or around *rpsT* or by increase of the gene copy number^13^. Similarly, mutations that disrupted mRNA structure of the *E. coli infA* gene, through local or long-range effects, explained much variation in fitness across a large collection of mutants^12^. Protein abundance and RNA structure contribute to the observed trans-effect of mutations^14^. Although our results are broadly consistent with a role of RNA structure, the specific structure is unknown, and the effects we uncovered are translation-independent, suggesting that a novel mechanism is involved. Toxic RNAs might interact with an essential cellular component, either nucleic acid or protein, and interfere with its normal function. Such interactions might be uncovered by pulldowns of toxic RNAs combined with sequencing or mass spectrometry. Alternatively, RNA phase transitions may be involved; such transitions have been shown to contribute to the pathogenicity of CAG-expansion disorders in Eukaryotes, providing a mechanistic explanation for this phenomenon^29^. Further studies will address the mechanisms, biotechnology applications, and evolutionary consequences of RNA toxicity in bacteria.

## Supplementary Methods

### Genes, plasmids and bacterial strains

We used a collection of 347 individually cloned full-length synonymous variants of the GFP gene. 154 of these variants came from our previous study^2^, while the others were ordered as gBlocks from Integrated DNA Technologies (IDT), generated from existing variants by DNA shuffling^19^ or made by site-directed mutagenesis, as described below. The coding sequences of all variants are provided as a fasta file. The GFP library was cloned into the Gateway entry plasmid pGK3, and then into Gateway destination plasmids pGK8, a T7 promoter-based expression plasmid, and pGK16, an expression plasmid with a bacterial *trp*/*lac* promoter^2^. Expression from both plasmids is IPTG-inducible; pGK8 produces untagged GFP, whereas pGK16 encodes a 28-codon 5′-terminal tag with weak mRNA secondary structure, known to facilitate expression^2^. pGK8 was used for GFP expression in the strain BL21-Gold(DE3) [*E. coli* B F^-^ *omp*T *hsd*S(r_B_^-^ m_B_^-^ *dcm*+ Tet^r^ *gal λ*(DE3) *end*A Hte], in C41/C43 strains (Lucigen)^22^, and in evolved suppressor strains (see below). pGK16 was used for expression in the DH5α strain [F^-^ Φ80*lac*Z Δ M15 Δ(*lac*ZYA-*arg*F) U169 *rec*A1 *end*A1 *hsd*R17(r_k_^-^, m_k_^+^) *pho*A *sup*E44 *thi*-1 *gyr*A96 *rel*A1 *λ*^-^]

### Growth assays

For growth rate analysis, three independent colonies of *E. coli* cells carrying each construct of GFP were grown overnight at 37°C in a 96-well plate with 2 ml wells, with constant shaking (320 rpm), until the cultures reached saturation. Following that, the culture was diluted 1: 100 in 200 μl of LB containing ampicillin (100 μg/ml) in a 96-well plate (Cat No. 655180, Cellstar). The plate was covered with its lid and placed in an automated plate reader (Tecan Infinite M200 Pro/ Tecan Sunrise). After an hour of incubating the plate at the appropriate temperature, typically 37°C, with constant orbital shaking (amplitude-1.5 mm, frequency-335 rpm), the cultures were induced with 1 mM IPTG. Subsequent to induction, plates were incubated in the plate reader with constant shaking as before. To avoid condensation while adding IPTG we retained the lid of the plate in the plate reader chamber which was maintained at 37°C and we avoided prolonged manipulation of plate outside the plate reader. To avoid excessive evaporation of cultures and test for potential contamination, we placed media without bacterial cultures in the external rows and columns of each plate, and only used internal wells for experiments. Optical density (OD) was measured at 595 nm and GFP fluorescence was measured with excitation at 485 nm and emission at 515 nm, at fixed time intervals over a period of 6-8 hrs (or 24 hrs in the low-temperature growth assays). LB-only wells were used to normalize the background OD and fluorescence. Bacterial growth rate and fluorescence represent means from three independent experiments, with three replicate measurements in each experiment.

We calculated the growth rates of IPTG-induced cultures as the slope of log2(OD) against time, normalized to the slope of non-induced cultures. Thus:

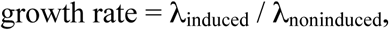
where λ = (log_2_(OD_t_) - log_2_(OD_0_)) / t; OD_0_ and OD_t_ represent the optical densities at the beginning and end of a time interval, and t represents the duration of the time interval. We defined the time interval as the interval between 1 h and 2.5 h after induction, with a further restriction that the OD of cells is between 0.1 and 1.0.

This formula gives a negative growth rate when the OD of the induced culture decreases over time, seen for example for GFP_170 in Figure 1B. We explain the slight reduction of OD by the lysis of a fraction of cells, mediated by expression of the toxic constructs. Indeed, we could observe cell lysis of GFP_170-expressing cells under the microscope.

Typically, the growth rate formula is applied to exponentially growing cells, and only gives positive values for such cells. In our experiments, although non-induced cells grew exponentially (Figure 1A), the growth rate of induced cells changed over time (Figure 1B), due to the combination of reduced growth following the expression of toxic constructs, partial cell lysis, and emergence of suppressors. Thus, the formula is only meant to approximate the behaviour of cells and provide a combined estimate of toxicity.

Graphs were plotted using GraphPad Prism 7 Software.

### Cell viability assay

The viability of bacterial cultures was estimated by spot assay or more quantitatively by measuring the colony forming units. For spot assays, BL21 strains carrying a subset of GFP constructs were grown in 2 ml LB with ampicillin, overnight in 14 ml falcon tubes with snap cap, at 37°C. Following growth to saturation, cells were diluted 1: 100 in LB containing ampicillin (100 μg/ml) and allowed to grow until OD reached ~0.5. Cultures in exponential phase were then diluted in LB (a factor of 10 between each step) and spotted on to LB plates containing Ampicillin (100 μg/ml) and 1 mM IPTG. Plate containing no IPTG were used as control to show equal number of cells were spotted. A volume of 10 μl was used for spotting. For quantitative measurements, exponential phase cultures were induced with 1 mM IPTG. Following induction, 1 ml of culture was aliquoted at every 1 hr interval and appropriately diluted (depending on OD) and 100 μl of appropriate dilution was spread on LB-Agar plates containing ampicillin (100 μg/ml). For each culture two different dilutions were spread at each time point in duplicate. Plates were then incubated at 37°C until colonies appeared on them. Viability was assessed by counting the colony forming units (cfu/ml) from the plates.

### Microscopic analyses of viability

Microscopy slides were prepared as previously published^30^. Briefly, two plain microscopy slides were cleaned with absolute ethanol. One of the two plastic covers of a gene frame (ABgene; 10 mm × 10 mm) was removed and the adhesive side pressed onto the centre of a glass slide. 1.5% Low Melting Point (LMP) agarose was dissolved in MQ water. 60 µl of the warm agarose solution was pipetted into the centre of each gene frame. The second glass slide was placed on top of the gene frame, avoiding the formation of any air bubbles. The sandwiched slides were allowed to set at 4°C for one hour. Then the upper glass slide was removed by sliding off gently from the agarose bed. BL21 cultures were grown in LB with ampicillin until OD reached ~0.2-0.3, following which they were induced with 1 mM IPTG. Un-induced cultures served as control. Subsequent to induction, aliquots were taken at appropriate time points. 1 µl propidium iodide (Life Technologies, 1mg/ml solution), that stains dead cells preferentially, was added to the aliquots. The tubes were then incubated at room temperature in dark for 5 minutes. 4 µl of culture was mounted onto the agarose bed and evenly spread on the agarose bed by turning the slide up and down. A clean glass coverslip was adhered to the upper adhesive side of the gene frame avoiding any air bubbles. Slides were imaged using using 100X Lens on a Zeiss Axio-Observer Z1 inverted microscope (Carl Zeiss UK, Cambridge, UK), with a ASI MS-2000 XY stage (Applied Scientific Instrumentation, Eugene, OR). Samples were illuminated using brightfield or a Lumencor Spectra X LED light source (Lumencor Inc, Beaverton, OR) complete with Chroma #89000ET single excitation and emission filters (Chroma Technology Corp., Rockingham, VT) and acquired on an Evolve EMCCD camera (Photometics, Photometrics, Tucson, AZ). GFP and RFP channels were used to image GFP and Propidium iodide (GFP-excitation: 470/22 nm, dichroic: 495 nm emission: 520/28 nm, RFP- excitation: 542/33 nm, dichroic: 562 nm emission: 593/40 nm). Image was captured using Micromanager (https://open-imaging.com/). For each microscopy slide, at least 10 independent fields were imaged in multi-channel acquisition mode, whilst remaining as unbiased as possible in order to obtain a true representation of the cell number and morphology of cells in the culture. Acquired images were analysed using ImageJ software.

### Generation of additional mutated constructs

To prevent the ribosomes from translating, we mutated the Shine-Dalgarno (SD) sequence in seven GFP constructs. All mutations were performed in the pGK3 plasmid^2^, using a site directed mutagenesis protocol^31^, employing AccuPrime™ Pfx DNA Polymerase (Thermo Fisher Scientific). The RBS site aaGAAGGA was changed to tgTTCTCT. The oligos used were SD_mut_Forward and SD_mut_Reverse primers (see List of oligos). The mutations were confirmed by sequencing and the constructs were then sub-cloned into pGK8 using Gateway cloning. The constructs were then transformed into BL21 cells and growth rates and fluorescence were analysed as described above.

Constructs expressing: 1) 132 nt fragments of GFP_012 and GFP_170 with and without start and stop codons (“Frag” and “Frag_(s+s)”), 2) GFP_012 and GFP_0170 with stop codons at 136^th^ and 157^th^ codon (“Stop1” and “Stop2”), and 3) GFP_012 and GFP_170 with transcription terminator sequence inserted at 492 nt position (“TT”), were generated as follows: 132 nt of GFP_170 and its corresponding region on GFP_012 were PCR-amplified using oligos containing BamHI and EcoRI sites (see List of oligos), for cloning into pGK3 plasmid. Start and stop codons were also added to the respective oligos in case of Frag_(s+s) constructs. To introduce TAA stop codons at 136^th^ and 157^th^ codon positions, site directed mutagenesis was carried out using specific oligos on pGK3-GFP_012 and pGK3-GFP_170 plasmids. In the same way we introduced stop codons in all three reading frames at the 157^th^ and 215^th^ codon positions of plasmid pGK3-GFP_170. To introduce Transcription Terminator (TT), 5′end phosphorylated oligos containing 57 nt sequence of TT, were self-annealed and cloned into the HpaI site of GFP_012 and GFP_170, on pGK3 plasmid. To fuse FLAG tags with the toxic fragment of GFP_170 (514-642bp) in all three reading frames, we amplified the GFP fragment from pGK3-GFP_170 with a forward primer containing a BamHI site and three individual reverse primers containing the FLAG tag in three different reading frames along with an EcoRI site. All the above constructs were cloned into pGK3, confirmed by sequencing and subcloned into pGK8 by Gateway cloning. All pGK8 constructs were then transformed into the BL21 strain and growth and fluorescence were analysed as above.

DNA shuffling was performed as previously reported^19^, with minor modifications. Briefly, an incomplete DNase I digestion of equimolar concentrations of the two variants was carried out in the presence of 5 mM MnCl_2_. Mn2+ ions in the reaction ensure DNaseI digests both strands of DNA at approximately the same sites^32^. To achieve controlled digestion DNase I treatment was performed for only 2 minutes at 15°C before inactivating the enzyme at 90°C for 5 minutes. Digested products were assembled by primerless assembly to obtain larger fragments of expected size. Assembly PCR was performed using Q5 high fidelity DNA polymerase (NEB) and PCR conditions were as follows: Annealing temperature:45°C, extension time: 30 secs for 40 cycles. The above step was followed by re-amplification with oligos pENTR_seq_U6 and pENTR_seq_L3. We obtained 36 GFP constructs from this experiment that were made of randomly shuffled fragments of GFP_012 and GFP_170. The shuffled variants encoding the GFP protein sequence were cloned into the pGK16 vector using Gateway cloning and transformed into DH5α for analysis of growth phenotype. To make synonymous mutations in the region spanning nts 534-642 in GFP_170, we designed degenerate oligos in five windows of 20-25 base pairs. In each window all wobble positions were mutated synonymously, allowing all possible changes at a given position. Site directed mutagenesis was performed using oligo sets A, B, C, D and E (see list of oligos) and AccuPrime Pfx DNA Polymerase (Thermo Fisher Scientific). All mutagenesis were carried out on the pGK3-GFP_170 plasmid. Mutations were confirmed by sequencing and the constructs were then sub-cloned into pGK8. The number of mutations per construct that we generated ranged from 2-9 and we obtained 98 constructs from five sets of PCRs. Single mutations were also generated in the region spanning 540-620 bp. Each wobble position was mutated synonymously, allowing all possible changes. We generated 36 constructs such that each construct had only one synonymous mutation per construct at a given codon in the region. Codons which were exactly the same between GFP_012 and GFP_170 were not mutated.

Bovine mitochondrial 2-oxoglutarate carrier protein (OGCP) constructs: wild type OGCP (OGCP_WT), OGCP with Shine-Dalgarno sequence changed from GAAGGA to TTCTCT and with no start codon (OGCP_noRBS), OGCP with *E. coli*-optimized codons (OGCP_CO), were purchased as gBlocks from IDT.

mKate2 constructs: A mKate2 gene fusion with the toxic fragment of GFP_170 was also ordered as a gBlock from IDT. The fragment contained BamHI and EcoRI sites for cloning into the pGK3 plasmid. The mKate2 gene by itself was amplified from the mKate-GFP_170 fusion construct using primers containing BamHI and EcoRI sites for cloning into pGK3. All constructs were confirmed by sequencing and subcloned into pGK8 by Gateway cloning.

### Isolation and validation of genetic suppressors

BL21 cells carrying several GFP variants (both toxic and non-toxic) were plated on LB agar supplemented with Ampicillin (100 μg/ml) and 1 mM IPTG. We obtained two kinds of colonies on the plates: highly fluorescent small colonies and large white colonies. We picked primarily the green colonies and a few white colonies for further analyses. All the colonies that were picked were plated on LB Agar+Amp plates. All green and some white colonies grew on Amp plates while some of the whites couldn’t grow any further on Amp plates. 37 colonies that grew on Amp plates (30 green and 7 white) were selected for further study. The growth rate and GFP fluorescence levels were measured for all colonies as described above.

To validate that the mutation that affects the survival of cells on IPTG is located on the chromosome and not on the plasmid itself, we cured the strains of the plasmid. For curing, the colonies were streaked on LB Agar plates in absence of Ampicillin repeatedly for at least 3-4 rounds. Colonies obtained after growing without antibiotic selection were further replica plated on LB Agar and LB Agar+Amp. Colonies that grew on only LB Agar but not on LB Agar+Amp were cured of the plasmid. These cured strains were re-transformed with the same GFP variants from which they were isolated. After re-transformation these cured strains were plated on LB Agar+Amp+IPTG plates. We obtained only bright green colonies from the re-transformed cured strain that was originally bright green. However, the cured strain from white colonies, on retransformation with the same GFP plasmid, produced a mix of green and white colonies on IPTG plates.

To further validate that the mutation was not located on the plasmid we isolated plasmids from the 37 colonies, and transformed them into fresh competent BL21 strain and assayed the growth and fluorescence. The phenotype was the same as in the parental strains, showing that the isolated plasmids did not carry any mutations that affected the phenotype. To identify the genomic mutations that conferred the suppressor phenotype we selected 22 suppressors (green= 18, white=4) for genome sequencing. We also sequenced the genomic DNA from two independent BL21 parental colonies to serve as reference and control during the analysis of genome sequences.

### Analysis of genome sequence and variant calling

Chromosomal DNA was isolated using the Wizard^®^ Genomic DNA Purification Kit (Promega, U.S.A.) according to the manufacturer’s instructions. The concentration of genomic DNA was estimated by Qubit dsDNA BR Assay Kit (ThermoFischer Scientific). Quantitation and quality control of genomic DNA was performed on a Bioanalyzer (Edinburgh Genomics UK). Genomic DNA samples were supplied in required concentrations for Nextera XT Library preparation, followed by 250-bp paired-end HiSeq Illumina sequencing (Edinburgh Genomics, UK).

The reads were mapped onto the reference genome sequence of BL21-Gold(DE3) (GenBank Accession ID CP001665.1) with default settings using bwa^33^. PCR duplicates were marked using Picard tools (http://broadinstitute.github.io/picard/). Genomic variants (SNPs, indels and insertions) were called using GATK^34^. We used GATK haplotype caller with ploidy= 1, stand_call_conf=30 and stand_emit_conf=10. Variants were filtered with parameter settings: DP<9.0 and QUAL<10.0. Bedtools^35^ was used to detect unique variation in our suppressor strains in comparison to the control strain and the reference genome. Finally, the identified variations were confirmed by targeted PCR amplification followed by Sanger sequencing. As the lac promoter is duplicated in the BL21-Gold(DE3) strain, wild type lac promoter (P*_lac_*_WT_) and the *lac*UV5 promoter(P*_lac_*_UV5_)driving the expression of T7 polymerase, we obtained dual peaks in targeted sequencing of P*_lac_*UV5 promoter region. To resolve this we carried out a detailed analysis of this region by extracting all read pairs where one read of the pair was mapped on to an unduplicated region and the read pairs were unambiguously assigned to the specific loci on the genome. The genome sequencing results can be accessed on https://www.ncbi.nlm.nih.gov/sra/SRP149903.

### Statistical analyses

We annotated the GFP sequences with a range of sequence-derived parameters and experimental measurements. The codon adaptation index (CAI) was calculated as in^2^ using codon optimality scores from^36^. GC3 content (GC content at the third positions of codons) was calculated by dividing the number of G- and C-ending codons by the total number of codons. Folding energy within the window (−4 to +38) relative to the translation start codon^2^ was calculated using hybrid-ss-min from the UNAFold package^37^. Translation initiation rate was calculated in a window from −40 to +60 relative to the start codon using the RBS Calculator^18^. Growth rate was calculated as described in the “growth assays” section above. OD was measured 3 hr after IPTG induction, after subtraction of LB-only background. OD and fluorescence measurements from a previous study^2^ were used after converting their units to units measured in the present study with a linear least squares model. Protein level measurements by Coomassie staining and RNA measurements by Northern blotting were from a previous study^2^. Protein abundance per cell was calculated by dividing protein fluorescence by OD.

To map the toxicity-determining region of GFP_170 based on the DNA shuffling experiment, we used Student’s t-test for each synonymous position *i* to compare the growth rates of variants in which position *i* was derived from GFP_170 and from GFP_012. We applied a Bonferroni correction for 239 tests, resulting in a p-value cutoff of 0.0002 (0.05/239). In this analysis, positions 532-640 from GFP_170 were associated with significantly slower growth of shuffled variants. We conservatively defined a slightly larger fragment (nts 512-645 of GFP_170) as the putative toxicity-determining region. We subsequently narrowed down this region based on the results of mutagenesis experiments. Regression analyses were performed in the R software package. Correlations reported in the text are quantified by the Spearman rank correlation coefficient and its associated p-value. We performed multiple regression analyses in order to quantify the relative importance of the various predictor variables in determining growth rates and optical densities. The output of these analyses, shown below, highlights the predominant influence of toxic mRNA fragments on growth:

Multiple regression. Dependent variable: growth rate, BL21 cells

| Coefficients | Estimate | Std. Error | t value | Pr(>\|t\|) | Significance |
| --- | --- | --- | --- | --- | --- |
| GFP_155_nt490-720 | -0.30111 | 0.05081 | -5.927 | 1.48E-07 | *** |
| GFP_170_nt514-645 | -0.41221 | 0.04715 | -8.743 | 2.05E-12 | *** |
| CAI | 0.24554 | 0.1678 | 1.463 | 0.148444 |  |
| GC3 | -0.03029 | 0.09525 | -0.318 | 0.751524 |  |

Signif. codes: 0 '\*\*\*' 0.001 '\*\*' 0.01 '\*' 0.05 '.' 0.1 ' ' 1
Residual standard error: 0.1077 on 62 degrees of freedom
Multiple R-squared: 0.6612, Adjusted R-squared: 0.6394
F-statistic: 30.25 on 4 and 62 DF, p-value: 5.752e-14

Multiple regression. Dependent variable: growth rate, DH5α cells

| Coefficients | Estimate | Std. Error | t value | Pr(>\|t\|) | Significance |
| --- | --- | --- | --- | --- | --- |
| GFP_155_nt490-720 | -0.19791 | 0.05044 | -3.924 | 0.000183 | *** |
| GFP_170_nt514-645 | -0.16322 | 0.03054 | -5.344 | 8.37E-07 | *** |
| CAI | 0.17294 | 0.17654 | 0.98 | 0.330226 |  |
| GC3 | -0.09352 | 0.08833 | -1.059 | 0.292866 |  |

Signif. codes: 0 '\*\*\*' 0.001 '\*\*' 0.01 '\*' 0.05 '.' 0.1 ' ' 1
Residual standard error: 0.1071 on 80 degrees of freedom
Multiple R-squared: 0.3628, Adjusted R-squared: 0.331
F-statistic: 11.39 on 4 and 80 DF, p-value: 2.296e-07

Multiple regression. Dependent variable: OD, BL21 cells

| Coefficients | Estimate | Std. Error | t value | Pr(>\|t\|) | Significance |
| --- | --- | --- | --- | --- | --- |
| GFP_155_nt490-720 | -0.20718 | 0.02904 | -7.135 | 2.29E-11 | *** |
| GFP_170_nt514-645 | -0.40684 | 0.03874 | -10.501 | < 2e-16 | *** |
| CAI | 0.36495 | 0.08492 | 4.298 | 2.82E-05 | *** |
| GC3 | -0.04033 | 0.06678 | -0.604 | 0.547 |  |

Signif. codes: 0 '\*\*\*' 0.001 '\*\*' 0.01 '\*' 0.05 '.' 0.1 ' ' 1
Residual standard error: 0.09221 on 180 degrees of freedom
Multiple R-squared: 0.5559, Adjusted R-squared: 0.546
F-statistic: 56.32 on 4 and 180 DF, p-value: < 2.2e-16

### Western blotting

For Western blotting, BL21strains carrying non-toxic and toxic GFP constructs +/- RBS were grown in 2 ml LB with ampicillin, overnight in snap cap tubes, at 37°C. Following growth to saturation, cells were diluted 1: 100 in LB containing ampicillin (100 µg/ml) and allowed to grow until OD reached ~0.5 and then induced with 1mM IPTG. Un-induced samples were collected before adding IPTG as control. After 1.5 h of induction, 1-2 ml of cultures were pelleted. Pellets were re-suspended in standard RIPA buffer and briefly sonicated in presence of Protease inhibitor (Roche) to lyse the cells. The lysate was further spun at 14000rpm for 5 mins to get rid of debris and the total protein was estimated by BCA assay (Pierce BCA protein estimation kit). 10µg of protein was resolved on 10% Bis-Tris gel. Prestained PageRuler protein ladder (ThermoFisher Scientific) was used as standard. Following electrophoresis the gel was transferred onto Nitrocellulose membrane using iblot2 gel transfer device (Invitrogen). The following antibodies were used for detection: Polyclonal Anti-GFP antibody (ab290, abcam), 1:5000, and goat anti-rabbit IgG-HRP conjugate (Santa Cruz Biotech, SC2030), 1:10000.

In the case of Flag fusion constructs, cells were grown and processed as described above. 13 µg of protein was resolved on 4-12% Bis-Tris gel. Prestained Benchmark protein ladder (Invitrogen) was used as standard. The following antibodies were used for detection, Flag M2 Monoclonal antibody (F3165, Sigma), 1:2000 and goat anti-mouse IgG-HRP conjugate (Santa Cruz Biotech, SC2031), 1:10000. The membranes were developed by soaking in Chemiluminescent substrate (Protein simple) and blots were imaged on Imagequant LAS4000 (GE Healthcare).

---

**Supplementary Figure 1.**
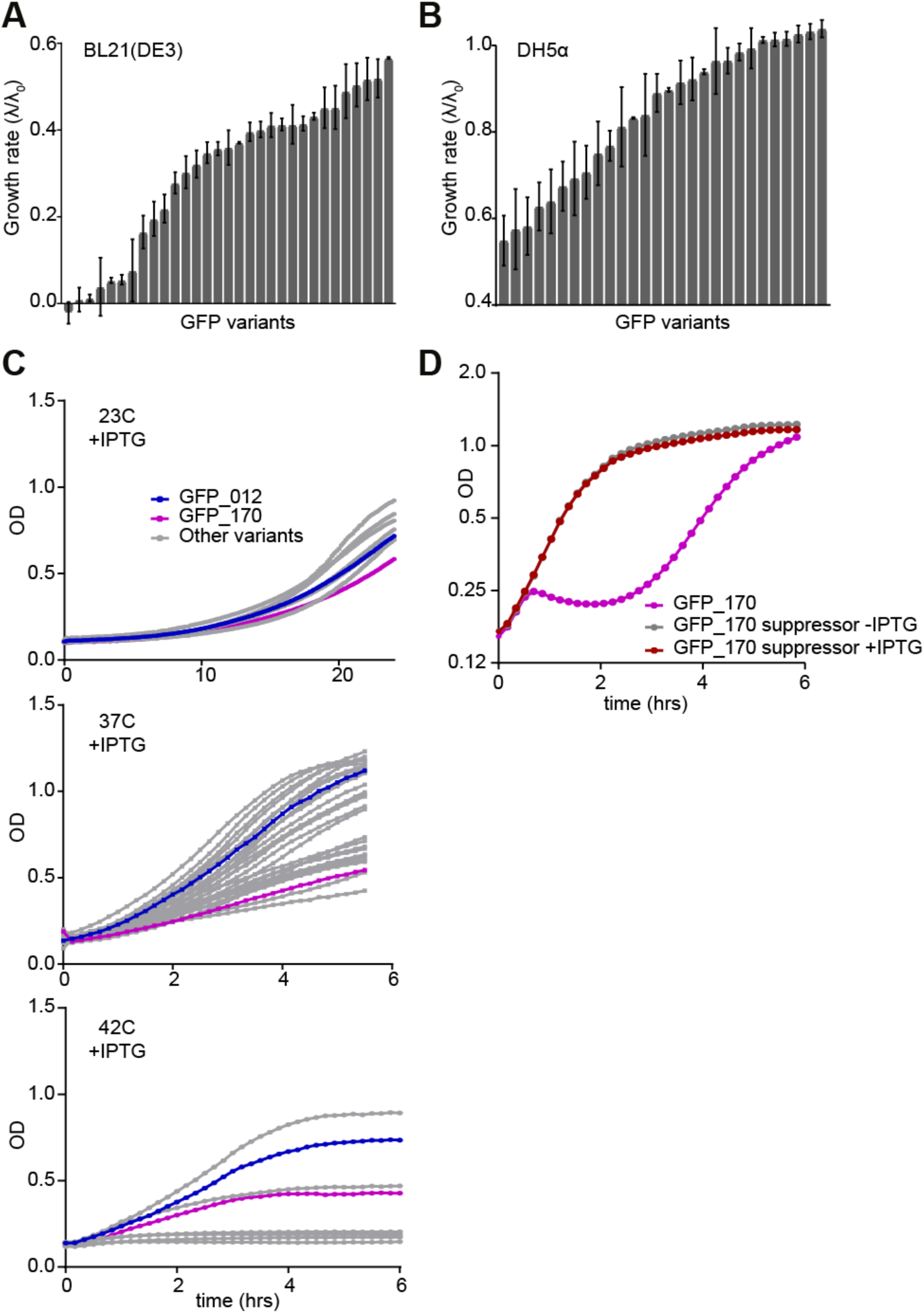
Growth phenotypes of GFP variants. **(A-B)** Growth rates in BL21 cells (A) and DH5α (B) in the presence of IPTG, sorted from minimum to maximum growth rate in each strain. **(C)** Growth curves of DH5α cells at different temperatures (23°C, 37°C and 42°C) in presence of IPTG. At 23°C there are minor variations in growth of cells expressing GFP variants, at 37°C there are large variations, and at 42°C, some of the GFP variants fail to grow altogether. GFP_012 (non-toxic, blue), GFP_170 (toxic, magenta), other variants (grey). The growth curves represent averages of at least 6 replicates. (D) Growth curve of BL21 cells expressing GFP_170 (magenta); suppressor isolated after back-diluting cells expressing GFP_170 in presence (red) and absence (grey) of IPTG. The suppressor strain has similar growth phenotypes both in presence and absence of IPTG.

**Supplementary Figure 2.**
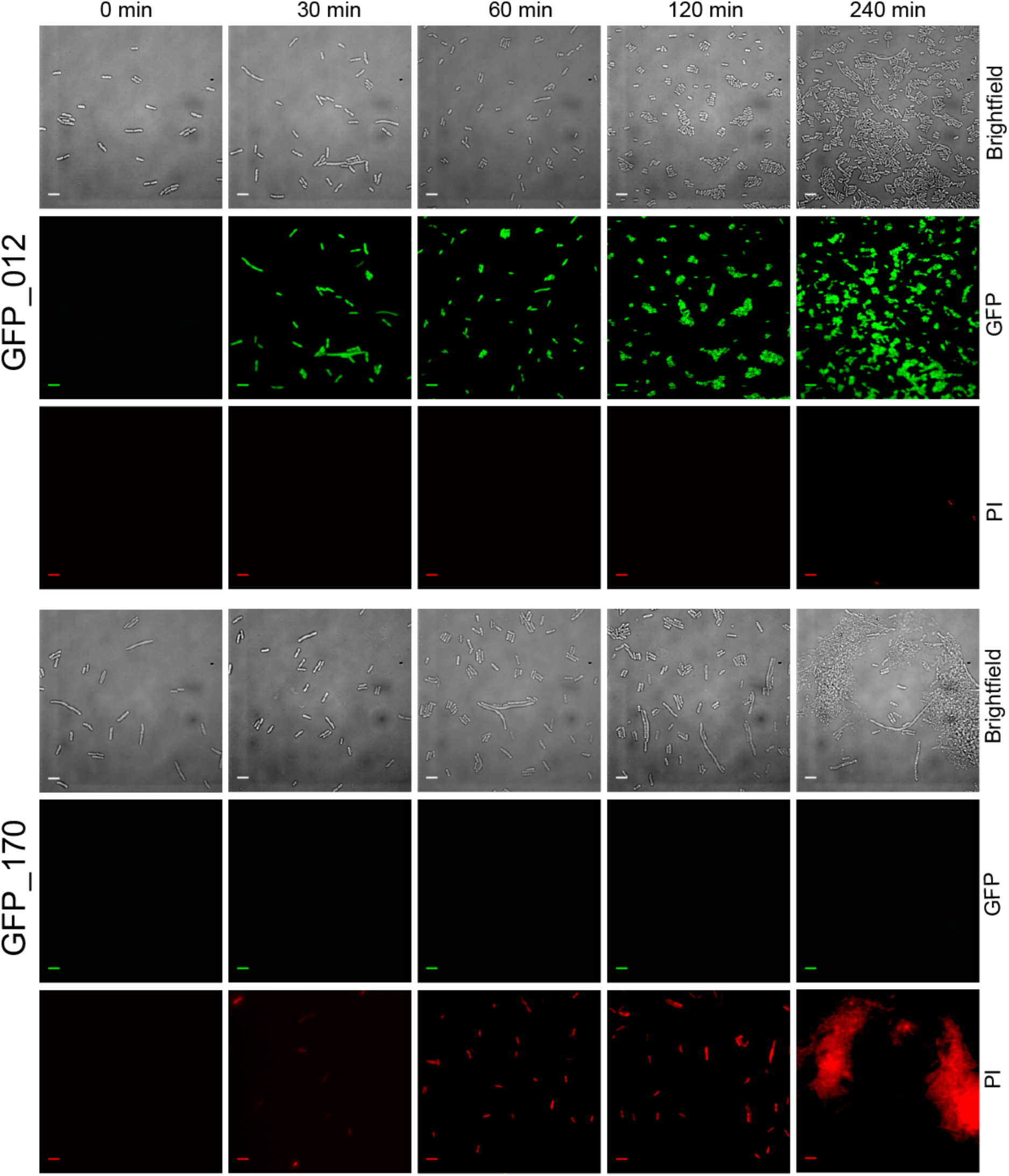
Microscopic analysis of cell viability. Cell viability was estimated for BL21 cells expressing GFP_012 (non-toxic variant) and GFP_170 (toxic variant). Brightfield images give an estimate of cell morphology and densities. GFP and RFP channels were used to determine the number of cells expressing GFP and the number of dead cells stained by Propidium Iodide (PI) respectively. At 0 min (just before IPTG induction) GFP_012 and GFP_170 cultures have similar cell densities and morphology. For cells expressing GFP_012, we see a steady increase in cell number after induction and GFP expression appears after 30 mins of induction. There is no significant cell death (PI stained cells) at any given time point. For cells expressing GFP_170 cell densities do not increase rapidly and most cells lose their morphology. We see a rapid increase in number of dead cells and the severity of the phenotype can be estimated at 240 min time point when PI staining shows only dead cells or debris from the dead cells. GFP expression is not seen for GFP_170 due to a strong mRNA secondary structure at its 5′ end, impeding its translation. The scale bar is 5 µm.

**Supplementary Figure 3.**
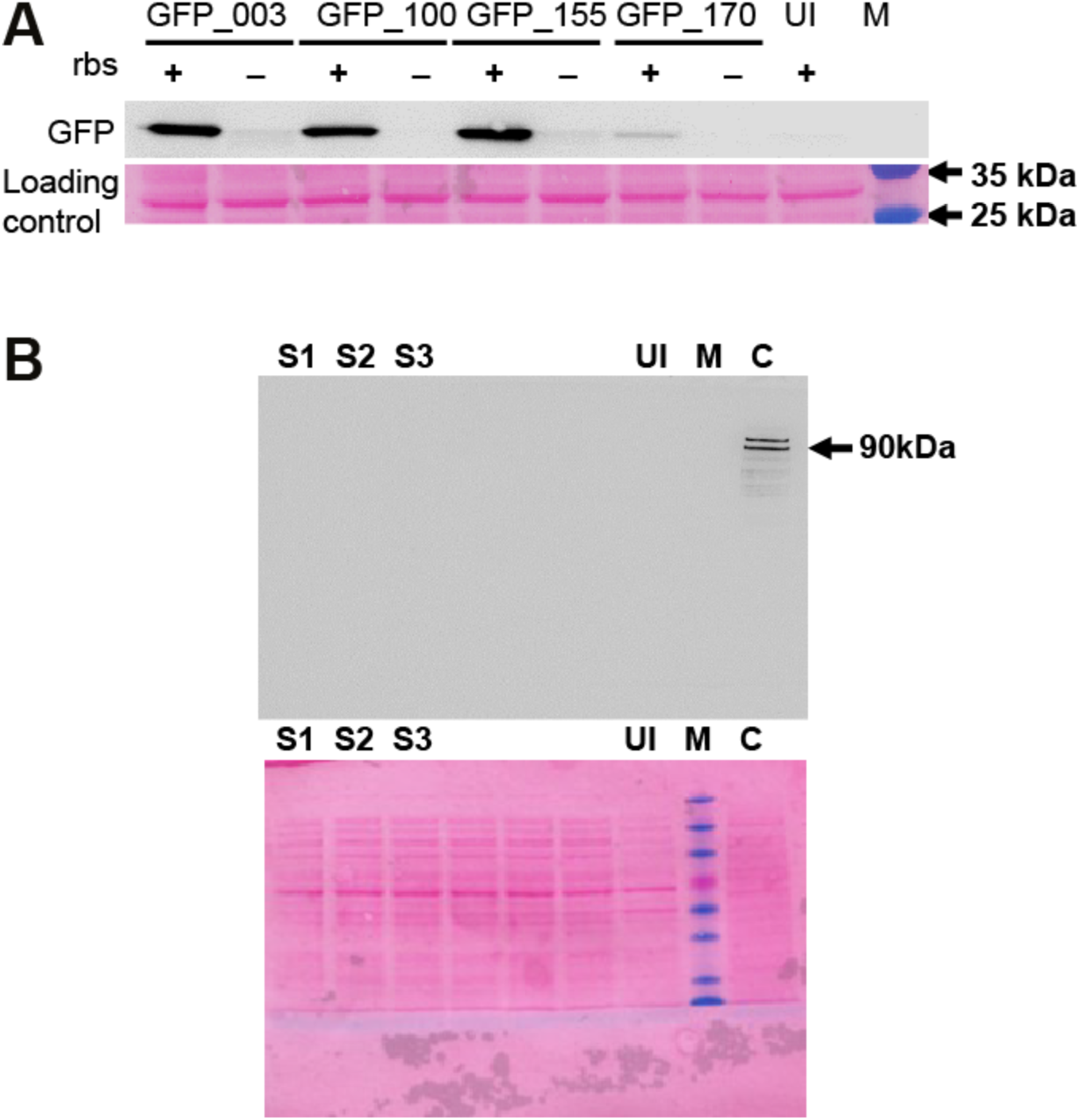
Measurement of GFP expression by Western blotting. **(A)** Expression of four toxic variants of GFP in the presence and absence of RBS. UI, uninduced control; M, marker. GFP expression was analysed by probing with anti-GFP polyclonal antibody (abcam 290). Ponceau stained blot shows equal loading. **(B)** GFP_170 toxic fragment (nt 514-645) expression fused to FLAG tag in all three reading frames (S1, S2, and S3) was analysed by probing with monoclonal Anti-FLAG (F3165 sigma). UI, uninduced control; M, marker; C, control sample expressing two Flag-tagged proteins of size 116 and 90 kDa. No FLAG expression was detected from S1, S2 or S3 constructs.

**Supplementary Figure 4.**
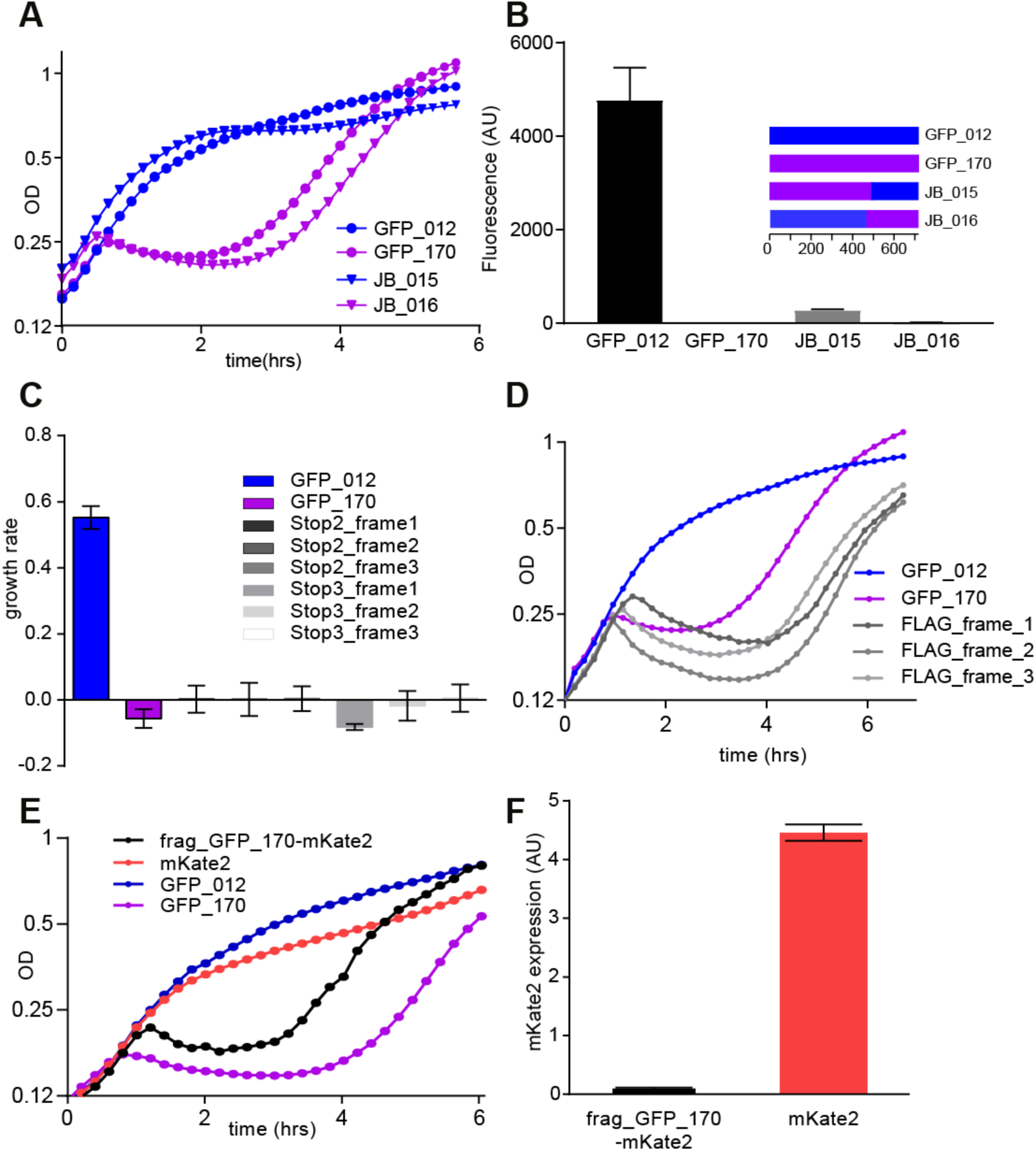
The toxic element resides near the 3′ end of GFP_170 and toxicity is independent of translation. **(A)** Growth curve for BL21 cells expressing constructs GFP_012, GFP_170 and their shuffled variants JB_015 and JB_016. JB_015 consists of GFP_170 (nts 1-497) and GFP_012 (498-720); JB_016 consists of GFP_012 (1-449) and GFP_170 (450-720). **(B)** Fluorescence of the shuffled constructs. JB_015 is non-toxic and shows a low level of fluorescence; JB_016 and GFP_170 are toxic and almost non-fluorescent. **(C)** Growth rate of cells expressing GFP_170 constructs with internal stop codons before and after the toxic fragment (nt 514-645) in all three reading frames. TAA stop codons were inserted at nucleotide positions 469 (stop2_frame1), 470 (stop2_frame2) and 471 (stop2_frame3) upstream of the toxic fragment and 643 (stop3_frame1), 644 (stop3_frame2) and 645 (stop3_frame3) downstream of toxic fragment. (D) Growth curves of constructs having toxic fragment from GFP_170 fused to FLAG tag at the 3′ end in all three reading frames. All three constructs retain toxicity. (E) Growth curves of mKate2 and toxic GFP_170 fragment fused to mKate2 at the 5′ end. Fusion construct retains toxicity (F) Expression of mKate2. No fluorescence is detected when mKate2 is fused with the toxic fragment from GFP_170.

**Supplementary Figure 5.**
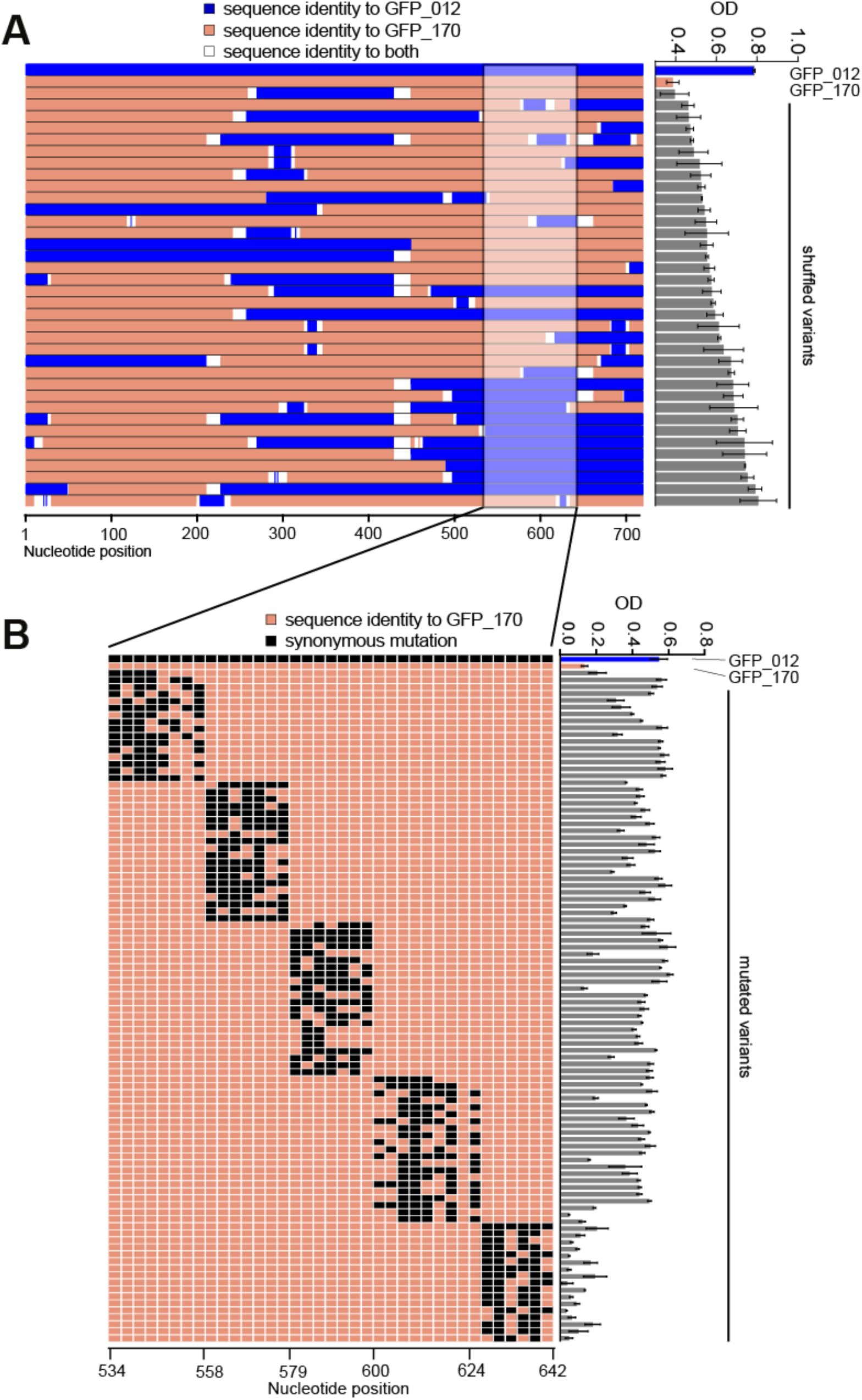
Growth analysis of GFP constructs generated by shuffling and multiple synonymous mutations. **(A)** 36 constructs were generated by DNA shuffling of GFP_012 (blue) and GFP_170 (orange). All constructs encode full length GFP. Constructs are colour coded according to the sequence identity with GFP_012 and GFP_170. The constructs from top to bottom are arranged in ascending order of their growth (OD 595nm). The highlighted region shows that most constructs having sequence identical to GFP_170 (orange) in 520-620 nt region are toxic. **(B)** An inset of the highlighted area from Panel A summarizes the results of multiple synonymous mutations that were generated in the toxic region. Each row represents a particular mutated variant and each column represents the nucleotide position. Columns highlighted orange and black represent nucleotides identical to GFP_170 and synonymous substitutions respectively. Each construct has 2-9 substitutions. Synonymous mutations in the region 534-624 nt reduce or abolish the toxicity of GFP_170 but any number of synonymous mutations in 627-642 nt region had no effect on toxicity. All data are averages of 9 replicates, +/- SEM.

**Supplementary Figure 6.**
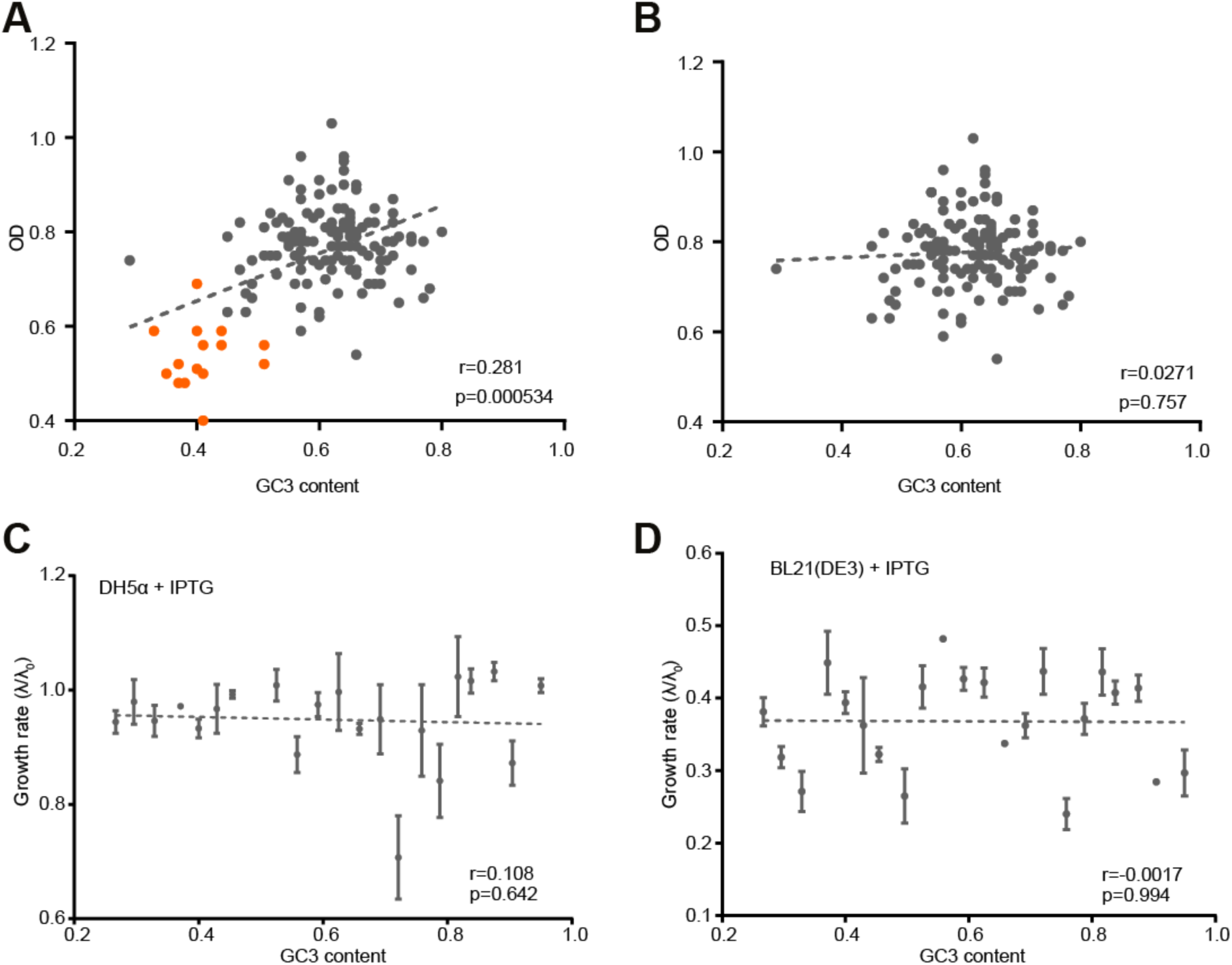
No correlation between GC3 content and growth rate of GFP variants. **(A-B)** The correlation between GC3 content and growth (OD 595nm) of GFP variants in BL21 cells is driven by two toxic RNA fragments shared between a number of variants: GFP_155 nt 490-720, and GFP_170 nt 514-645, marked in orange. After removal of these variants (panel B), we no longer see any relationship between GC3 content and growth. **(C-D)** There is no relationship between GC3 content and growth in an independent set of 22 GFP constructs, either in DH5α (C) or BL21 (D) strains. All data are averages of 9 replicates, +/- SEM.

**Supplementary Figure 7.**
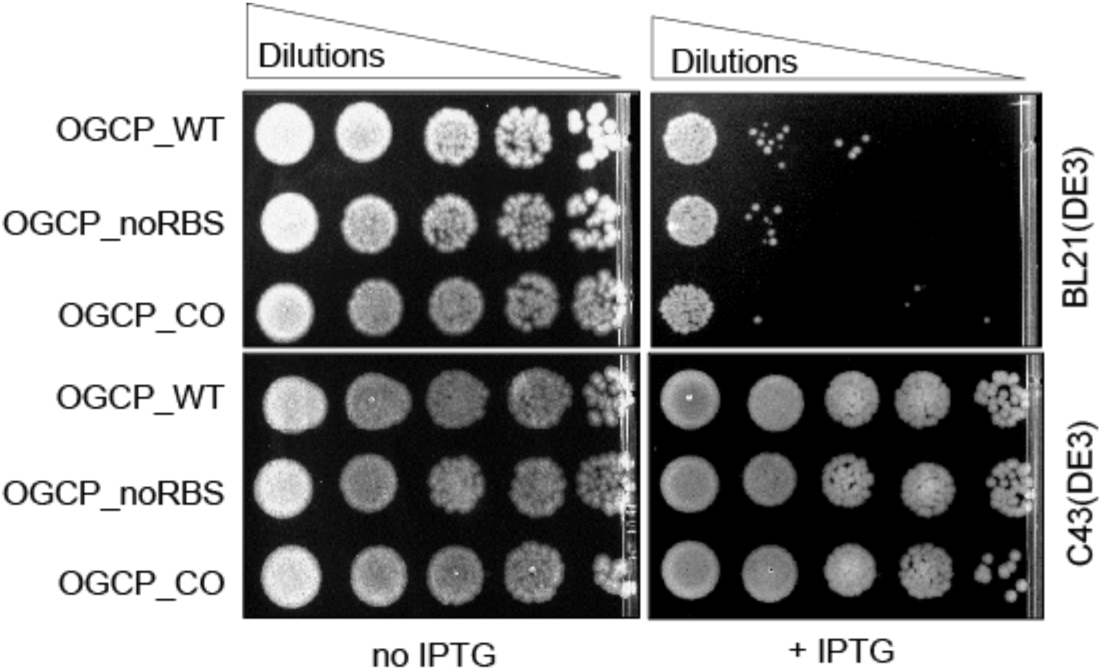
Spot assay for semi-quantitative estimation of cell viability of BL21 cells expressing OGCP variants. OGCP-WT (wild type OGCP), OGCP_noRBS (OGCP lacking functional RBS) and OGCP_CO (codon-optimized OGCP) variants were cloned in pGK8 plasmid and transformed in BL21 and C43 strains. In the absence of IPTG there are no difference in the viabilities between strains or constructs; in the presence of IPTG, the three constructs are toxic in BL21 cells but not in C43 cells.

**Supplementary Table 1.** Analysis of suppressor genotypes. 15/18 green suppressors showed a complete replacement of P*_lac_*_UV5_ promoter with P*_lac_*_WT,_ 3/18 showed replacement of P*_lac_*_UV5_ with P_l*ac*Weak._ 3/4 white suppressors had no changes in the promoter of T7 RNA polymerase, while for 1/4 we could not definitively assign the promoter type.

| Strain | Isolated from | Color | Genotype |
| --- | --- | --- | --- |
| Sup_01 | GFP_003 | Green | P<sub>lacWeak</sub> |
| Sup_02 | GFP_003 | Green | P<sub>lacWT</sub> |
| Sup_03 | GFP_003 | Green | P<sub>lacWeak</sub> |
| Sup_04 | GFP_003 | Green | P<sub>lacWT</sub> |
| Sup_05 | GFP_003 | Green | P<sub>lacWT</sub> |
| Sup_06 | GFP_003 | Green | P<sub>lacWT</sub> |
| Sup_07 | GFP_003 | Green | P<sub>lacWT</sub> |
| Sup_08 | GFP_003 | Green | P<sub>lacWT</sub> |
| Sup_10 | GFP_003 | Green | P<sub>lacWT</sub> |
| Sup_12 | GFP_003 | White | P<sub>lacUV5</sub> |
| Sup_14 | GFP_069 | Green | P<sub>lacWT</sub> |
| Sup_15 | GFP_069 | Green | P<sub>lacWT</sub> |
| Sup_17 | GFP_069 | Green | P<sub>lacWT</sub> |
| Sup_18 | GFP_069 | White | P<sub>lacUV5</sub> |
| Sup_21 | GFP_183 | Green | P<sub>lacWT</sub> |
| Sup_22 | GFP_183 | Green | P<sub>lacWeak</sub> |
| Sup_24 | GFP_183 | White | P<sub>lacUV5</sub> |
| Sup_26 | GFP_155 | Green | P<sub>lacWT</sub> |
| Sup_30 | GFP_155 | Green | P<sub>lacWT</sub> |
| Sup_34 | GFP_100 | Green | P<sub>lacWT</sub> |
| Sup_35 | GFP_100 | Green | P<sub>lacWT</sub> |
| Sup_37 | GFP_170 | White | P<sub>lacUV5</sub>/P<sub>lacWeak</sub>** |

\*\* We observed a mix of two types of reads in sequencing analysis for this strain

